# Cell-Tak coating may cause mis-normalization of Seahorse metabolic flux data

**DOI:** 10.1101/2021.03.19.436150

**Authors:** Michal Sima, Stanislava Martinkova, Anezka Kafkova, Jan Pala, Jan Trnka

## Abstract

Metabolic flux investigations of cells and tissue samples are a rapidly advancing tool in diverse research areas. Reliable methods of data normalization are crucial for an adequate interpretation of results and to avoid a misinterpretation of experiments and incorrect conclusions. The most common methods for metabolic flux data normalization are to cell number, DNA and protein. Data normalization may be affected by a variety of factors, such as density, healthy state, adherence efficiency, or proportional seeding of cells.

The mussel-derived adhesive Cell-Tak is often used to immobilize poorly adherent cells. Here we demonstrate that this coating may strongly affect the fluorescent detection of DNA leading to an incorrect and highly variable normalization of metabolic flux data. Protein assays are much less affected and cell counting can virtually completely remove the effect of the coating. Cell-Tak coating also affects cell shape in a cell line-specific manner and may change cellular metabolism.

Based on these observations we recommend cell counting as a gold standard normalization method for Seahorse metabolic flux measurements with protein content as a reasonable alternative.

## Introduction

The measurement of cellular metabolism is a widely used research approach in a variety of disciplines. Any interventions that lead to a change in the physiological functioning of cells e.g. mutations, chemical treatments, environmental conditions and others can affect cellular metabolism. The extracellular flux (XF) measurement technology developed by Seahorse (Agilent) is an elegant method of measuring oxygen consumption and extracellular acidification rates in relatively small amounts of live biological material. We have previously used the Seahorse analyzer to study the effect of lipophilic cations on mitochondrial metabolism [1] inhibitory effect of the lipophilic positively charged moiety of methyltriphenylphosphonium (TPMP) on 2-oxoglutarate dehydrogenase [2] and the effect of Cu(II)–phenanthroline complexes on cellular metabolism [3] and in other studies.

The XF data usually requires a normalization due to the varying number of cells in each tested well – this requirement is most needed for *ex vivo* samples or when different cell lines are used in one experiment or to compare experiments from various times. A range of normalization strategies for XF metabolic assays are available such as normalization to total cellular protein [4], to nuclear DNA [5], to cell number calculated by microscope image analysis [6,7] or to the number of cell nuclei using fluorescent microscopy [8].

During measurements which include a degree of liquid agitation and mixing some cells tend to detach from the surface of the microplate wells, which can lead to unusable measurement data. It is thus common to coat the surface with agents that enhance the adhesion to the plastic [9]. Cells prefer to adhere to hydrophilic surfaces or surfaces that contain functional –NH_2_/–COOH groups [10,11].

One highly adhesive and a widely used coating material is Cell-Tak. Its main components are polyphenolic proteins extracted from the marine mussel *Mytilus edulis*, which has a remarkable ability to adhere to underwater surfaces [12–14]. Observations showed that these proteins rich in lysine, hydroxylated amino acids, and 3,4-dihydroxyphenylalanine have strong adhesive properties *in vitro* and contributes to byssal adhesion [12,15]. Multiple polyphenolic proteins were extracted from *M. edulis* and used as base of Cell-Tak [12,13]. Cell-Tak has been used for cell attachment to microscope slides in order to stabilize them for observation [16].

Some of our previous experiments using Cell-Tak for enhancing cellular adhesion showed inconsistent results of measured DNA concentrations used to normalize XF data. We investigated possible sources of this inconsistency and in this study we discuss the possible XF data normalization errors due to Cell-Tak coating. To exclude factors that affect measurement such as cell debris, protein, or lysis buffer content we started with DNA and protein standards with a known concentration on coated vs. non-coated surfaces and then followed with two different cell lines. Experiments were performed on standard plastic 96 well plates with four coating protocols and on a XF Seahorse plate with two coating protocols. Finally, we normalized acquired metabolic flux data based on the measured DNA assay fluorescence, protein assay absorbance and to the number of cells counted using a microscope. Our results indicate a clear effect of the used coating type on the normalized data which could lead to data misinterpretation.

## Material and Methods

### Cell lines, culture, and standards

Immortalized mammalian cell lines HepG2 (human liver cell line) and the C2C12 (mouse myoblast cell line) were kindly provided by Dr. Julien Prudent (MRC Mitochondrial Biology Unit, Cambridge, UK) and grown in Dulbecco’s Modified Eagle’s Medium (Life Technologies, cat. n. 31885023) supplemented with 10% FBS (Life Technologies, cat. n. A3160402) and 1% penicillin/streptomycin (Sigma-Aldrich, cat. n. P4333) at 37°C in 5% CO_2_. Cells were harvested using trypsin/EDTA (Life Technologies, cat. n. 15400054) and centrifuged at 150 x g for 5 minutes. Pellets were resuspended in complete Seahorse XF DMEM medium, pH 7.4 or in DMEM without phenol red (Life Technologies, cat. n. A1443001) and cells were counted under a Motic inverted microscope/AE20 microscope using a Bürker counting chamber.

As a DNA standard, the Lambda DNA in TE from Quant-iT ^™^ PicoGreen^®^ dsDNA Assay Kit (ThermoFisher Scientific, cat. n. P11496) was used. Bovine serum albumin (Sigma-Aldrich, cat. n. P0914-10AMP) was utilized as a protein standard.

The various treatments and assays are summarized in Table 1 and described below. The first series of experiments was done on 96 well plate with a DNA or protein standard with four plate coating protocols followed by analogous experimental settings but with two cell lines. After seeding and attachment cells were lysed and their DNA or protein content analyzed.

**Table 1.**
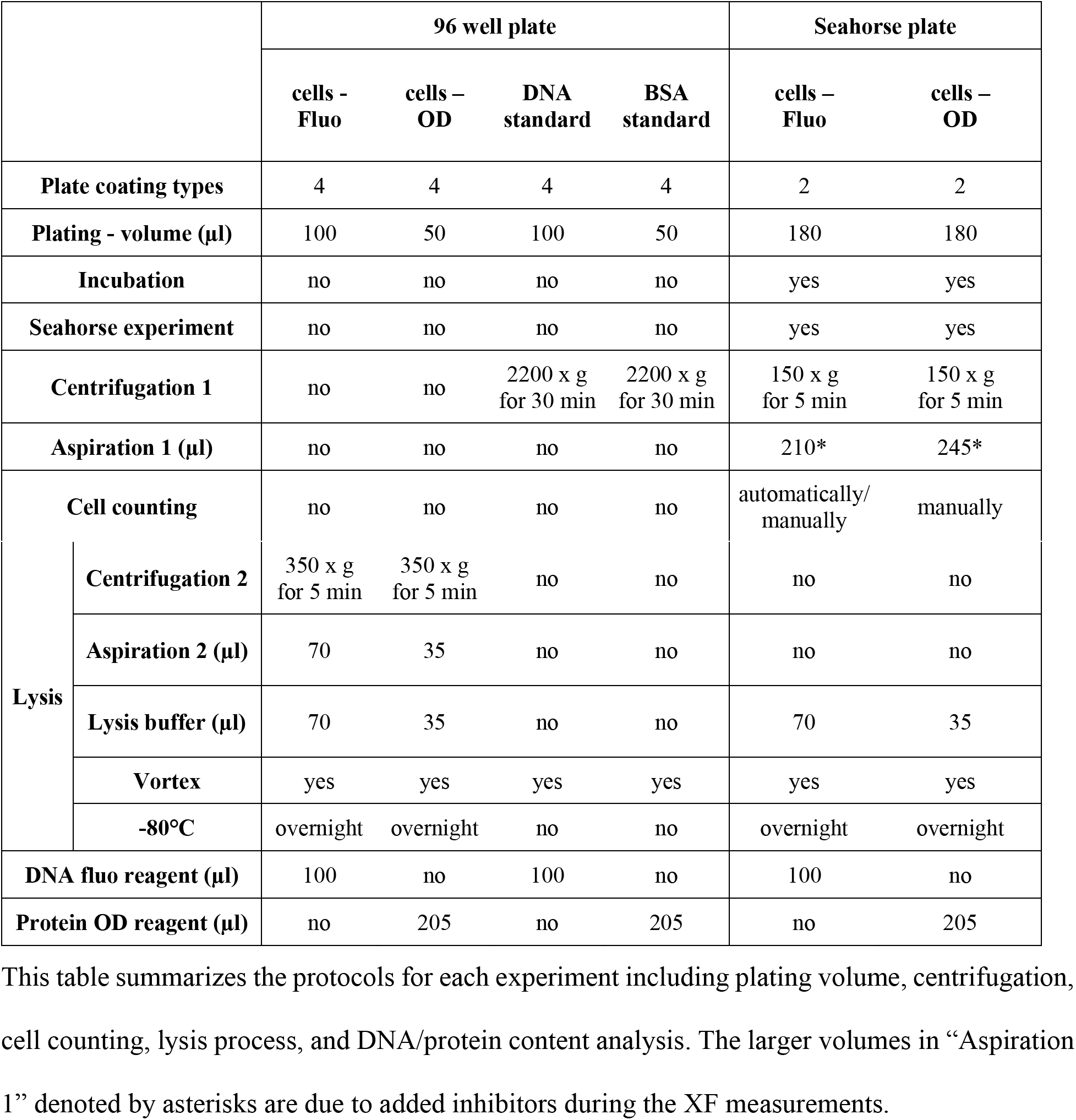
Summary of experimental protocols.

|  | 96 well plate |  |  |  | Seahorse plate |  |
| --- | --- | --- | --- | --- | --- | --- |
|  | cells -<br>Fluo | cells –<br>OD | DNA<br>standard | BSA<br>standard | cells –<br>Fluo | cells –<br>OD |
| **Plate coating types** | 4 | 4 | 4 | 4 | 2 | 2 |
| **Plating - volume (µl)** | 100 | 50 | 100 | 50 | 180 | 180 |
| **Incubation** | no | no | no | no | yes | yes |
| **Seahorse experiment** | no | no | no | no | yes | yes |
| **Centrifugation 1** | no | no | 2200 x g<br>for 30 min | 2200 x g<br>for 30 min | 150 x g<br>for 5 min | 150 x g<br>for 5 min |
| **Aspiration 1 (µl)** | no | no | no | no | 210* | 245* |
| **Cell counting** | no | no | no | no | automatically/<br>manually | manually |

|  |  |  |  |  |  |  |  |
| --- | --- | --- | --- | --- | --- | --- | --- |
| **Lysis** | **Centrifugation 2** | 350 x g<br>for 5 min | 350 x g<br>for 5 min | no | no | no | no |
|  | **Aspiration 2 (µl)** | 70 | 35 | no | no | no | no |
|  | **Lysis buffer (µl)** | 70 | 35 | no | no | 70 | 35 |
|  | **Vortex** | yes | yes | yes | yes | yes | yes |
|  | **-80°C** | overnight | overnight | no | no | overnight | overnight |
| **DNA fluo reagent (µl)** |  | 100 | no | 100 | no | 100 | no |
| **Protein OD reagent (µl)** |  | no | 205 | no | 205 | no | 205 |

This table summarizes the protocols for each experiment including plating volume, centrifugation, cell counting, lysis process, and DNA/protein content analysis. The larger volumes in “Aspiration 1” denoted by asterisks are due to added inhibitors during the XF measurements.

In the second type of experiments cells were seeded on XFp Seahorse plates with two types of coating. After the XF measurement sequence cells were counted and used for DNA content detection or counted and used for protein content analysis.

### Plating of standards and seeding of cells

For the first set of experiments we used the Nunc^™^ MicroWell^™^ 96-Well Microplates (ThermoFisher Scientific, cat. n. 269620) with Nunc^™^ Microplate Lids (ThermoFisher Scientific, cat. n. 263339). Four different variants of coating solutions were applied to these plates: 1) dH_2_O as a control, 2) 0.1 M NaHCO_3_ as another control, 3) Cell-Tak – Corning^®^ Cell-Tak^™^ Cell and Tissue Adhesive (Corning, cat. n. 354240) (3.5 μg/cm^2^) diluted in dH_2_O, and 4) Cell-Tak (3.5 μg/cm^2^) diluted in 0.1 M NaHCO_3_.

DNA and BSA standards were diluted in dH_2_O and added to the wells in concentrations 250 ng/ml and 500 ng/ml for DNA and 50 μg/ml and 100 μg/ml for BSA in volumes indicated in Table 1. Cells in DMEM without phenol red were added to the plates in two amounts: 10 000 and 20 000/well. Plates with standards were centrifuged at 2200 x g for 30 minutes and then vortexed briefly before the addition of DNA/protein analysis reagents (see below). Wells with dH_2_O only were used as a blank for standard analysis and wells with medium only served as a blank for cell analysis. All experiments were set up in triplicate and repeated three times.

The second set of experiments was performed on Seahorse XF Cell Culture Microplates (Agilent, cat. n. 103022-100) with eight wells. Cells were seeded on these plates as described above but only two versions of coating solutions were applied to these plates: four wells were coated with dH_2_O as a no-coating control, and the remaining four were coated with Cell-Tak (3.5 μg/cm^2^) diluted in 0.1 M NaHCO_3_ as per manufacturer’s instructions. Cells were seeded at 6000/well in Seahorse XF DMEM medium, pH 7.4 (Agilent, cat. n. 103575-100). The assay medium only was used as a blank in one well coated with dH_2_O and one well with Cell-Tak.

Both types of plates used in this study are made of hydrophobic untreated polystyrene with a flat bottom shape. All experiments were performed three times on separate days.

### XF measurements

Directly after seeding, cells were allowed to settle down for 20 minutes on the bench to promote an even distribution, and then transferred for 40 minutes to a 37°C/5% CO2 incubator. After that time preheated 37°C Seahorse XF DMEM assay medium, pH 7.4 was added to each well for a final volume of 180 μl/well. Medium was supplemented with 5.55 mM Glucose, 1 mM Sodium Pyruvate (Sigma-Aldrich, cat. n. S8636), and 4 mM L-Glutamine (Sigma-Aldrich, cat. n. G7513). Plates were placed in a non-CO2 incubator at 37°C (according to manufacturer’s instructions) prior to the assay. Cellular mitochondrial respiration (OCR – oxygen consumption rate) was determined using the XFp analyzer (Agilent Technologies, CA, USA). Mitochondrial stress assay was performed with the consecutive 20 μl injections/each of reagents (final concentration): oligomycin (1 μM), carbonyl cyanide-4-(trifluoromethoxy)phenylhydrazone (FCCP)(1 μM – HepG2; 2 μM – C2C12), and antimycin A/rotenone (1 μM each). The last injection added 0.2 μg/ml Hoechst (Life Technologies, cat. n. 33342-Invitrogen^™^) used for cell counting. A total of 12 OCR measurements were taken - three for the basal respiration and three after each inhibitor injection.

### Cell counting

After the Seahorse experiment the plates were centrifuged (5 minutes at 150 x g) to sediment detached cells and a part of the medium was aspirated from the wells (Table 1). Samples in Seahorse XF Cell Culture Microplates were thereafter scanned in the bright field mode by a monochrome fluorescence CCD camera Leica DFC 350FX mounted on camera port of inverted fully motorized microscope stand Leica DMI 6000. A Leica HC PL FLUOTAR 10x/0, 30 DRY objective was used to acquire tile scans with 4×5 fields with a corresponding pixel size 921×921 nm.

We then used the Fiji software [17] to count cells manually. During cell counting, cells were divided into two groups based on their shape - a round form, with defined and visible edges and flat - not round form, without defined edges or with protrusions.

### Cell lysis

Prior to DNA/protein content assay 96 well plates with cells were centrifuged for 5 minutes at 350 x g. Part of the medium from these plates was aspirated from the wells. Analogous steps were performed with Seahorse plates before cell counting (Table 1). Appropriate volumes of lysis buffer (Sigma-Aldrich, cat. n. C3228-500ML) were added to both plate types (Table 1). Plates were vortexed briefly and put for ten minutes at 37°C (three times repeated). To complete the lysis they were then placed into a −80°C freezer overnight.

### DNA and protein content assays

All measurements were performed in the original plates with standards/cells. For the determination of DNA content 100 μl of PicoGreen were added to 100 μl of DNA standard/thawed cell lysates. Fluorescence intensity was measured using TECAN Infinite M200Pro microplate reader (Schoeller instruments) with gain set manually to 80.

For the measurement of the total protein content 205 μL of Bradford Reagent (Sigma-Aldrich, cat. n. B6916) was pipetted to the 50 μL of BSA standard/thawed cell lysates. The absorbance was measured at wavelength 595 nm using TECAN Infinite M200Pro microplate reader.

### Data analysis

Three replicates in each experiment/treatment were averaged (fluorescence/optical density/cell number) and the appropriate blank averages (fluorescence/optical density) were subtracted from these values. We used wells “coated” with dH_2_O as the negative control in our experiment and all the values obtained from other coating options were normalized to this negative control them to prevent day-to-day signal variation. The resulting ratios of fluorescence/optical density/number of cells therefore indicate the effect of the coatings with respect to no coating (dH_2_O). These ratios from three independent experiments were then statistically analyzed.

The analysis of Seahorse data was performed as follows: for each measurement time-point, three measurement replicates were averaged to give the value for each experiment. Averages and standard deviations were then calculated from three independent experiments. These data were then normalized to the corresponding DNA/protein/cell number in the Wave data analysis software (Agilent) and exported to Microsoft Excel.

The measured values of fluorescence/absorbance/respiration for different experimental conditions taken from three independent experiments were compared using Student’s t-test and the respective p-values are denoted in figures and tables. The counts of cell shapes from three independent experiments were compared between treatments using Fisher’s exact test. We used GraphPad Prism 8 for the statistical analysis.

## Results

### Effect of coating on DNA and protein assays of standards

The first set of experiments was performed in 96 well plates with four different coating options: 1) dH_2_O, 2) 0.1 M NaHCO_3_, 3) Cell-Tak diluted in dH_2_O and 4) Cell-Tak diluted in 0.1 M NaHCO_3_, and with two standards in two concentrations: a) DNA (250 and 500 ng/ml) and b) BSA (50 and 100 μg/ml).

After plating into uncoated/coated wells (see Methods) they were analyzed by using PicoGreen (DNA content - fluorescence) and the Bradford reagent (protein content - absorbance), respectively.

Compared to the negative control (dH_2_O “coating”) there was no significant change in detected absorbance for both BSA concentrations and the data exhibited low standard deviations in all cases (Table 2). When DNA was used as a standard no significant fluorescence change was measured in wells coated with NaHCO_3_ compared to dH_2_O coating but an approximately 25% decrease of fluorescence was detected in wells coated with Cell-Tak diluted in dH_2_O and more than 50% decrease in wells coated with Cell-Tak diluted in NaHCO_3_ (Table 2) even if the amount of DNA was the same in all compared wells. These differences reached relatively low p-values. In the last two mentioned set ups large standard deviations indicate a high variability of the signal (Table 2).

**Table 2.** Mean change in DNA fluorescence and protein absorbance due to coatings.

|  |  |  |  | NaHCO<sub>3</sub> vs dH<sub>2</sub>O |  | (Cell-Tak in dH<sub>2</sub>O) vs dH<sub>2</sub>O |  | (Cell-Tak in NaHCO<sub>3</sub>) vs dH<sub>2</sub>O |  | NaHCO<sub>3</sub> vs Cell-Tak in NaHCO<sub>3</sub> |
| --- | --- | --- | --- | --- | --- | --- | --- | --- | --- | --- |
|  |  |  |  | aver | stdev | aver | stdev | aver | stdev | p-value |
| standards | FLUO | DNA (ng/mL) | 250 | 1,053 | 0,056 | 0,759 | 0,180 | 0,443 | 0,243 | 0,043 |
|  |  |  | 500 | 1,063 | 0,071 | 0,765 | 0,218 | 0,488 | 0,229 | 0,039 |
|  | OD | BSA (µg/mL) | 50 | 1,028 | 0,017 | 1,016 | 0,011 | 1,003 | 0,025 | 0,247 |
|  |  |  | 100 | 0,994 | 0,010 | 0,992 | 0,023 | 0,958 | 0,023 | 0,101 |
| cells | FLUO | HepG2/ well | 10000 | 1,341 | 0,073 | 1,327 | 0,077 | 1,097 | 0,133 | 0,066 |
|  |  |  | 20000 | 1,389 | 0,159 | 1,365 | 0,138 | 1,180 | 0,216 | 0,254 |
|  |  | C2C12/ well | 10000 | 1,133 | 0,063 | 1,115 | 0,112 | 0,655 | 0,040 | 0,001 |
|  |  |  | 20000 | 1,081 | 0,031 | 1,064 | 0,046 | 0,915 | 0,106 | 0,105 |
|  | OD | HepG2/ well | 10000 | 1,186 | 0,189 | 1,028 | 0,047 | 0,890 | 0,063 | 0,102 |
|  |  |  | 20000 | 1,065 | 0,040 | 1,023 | 0,017 | 0,898 | 0,044 | 0,008 |
|  |  | C2C12/ well | 10000 | 1,055 | 0,001 | 1,063 | 0,027 | 0,846 | 0,023 | 0,004 |
|  |  |  | 20000 | 1,017 | 0,331 | 0,990 | 0,383 | 0,875 | 0,369 | 0,647 |

Changes in DNA fluorescence (FLUO) and protein absorbance (OD) between the various coatings compared to dH<sub>2</sub>O control. Experiments with DNA and protein standards and with the two cells lines are shown here. The last column indicates the p-values from Student's t-test, when the ratios from NaHCO<sub>3</sub> vs dH<sub>2</sub>O and (Cell-Tak in NaHCO<sub>3</sub>) vs dH<sub>2</sub>O were compared among three independent experiments.

### Coating effects on DNA and protein assays on cultured cells

We performed an analogous set of experiments with the four coating options as above and with two cell lines: 1) HepG2 and 2) C2C12, at two densities: 10 000 and 20 000 cells/well. After cell lysis we measured DNA fluorescence or protein absorbance as before.

When assayed for protein content no significant differences were found in cells growing in wells coated with NaHCO_3_ and Cell-Tak/dH_2_O compared to dH_2_O (Table 2). In wells coated with Cell-Tak/NaHCO_3_ we observed a 10-15% decrease of absorbance (Table 2). When assayed for DNA content there we observed an increase in wells with HepG2 cells when coated with NaHCO_3_ and Cell-Tak/dH_2_O (approximately 35%) and a similar but smaller increase in C2C12 cells (6-10%) (Table 2). In wells coated with Cell-Tak/NaHCO_3_ a small increase of fluorescence was detected in HepG2 cells but fluorescence decreased in C2C12 cells (Table 2).

We then performed a similar set of experiments in the multi-well plates used for the metabolic flux measurements in the Seahorse machine and used cell counting in wells as an independent normalization method. There was a similar decrease of the DNA fluorescence for both cell lines (Table 3) in wells coated with Cell-Tak/NaHCO_3_ compared dH_2_O (approximately 15%). Protein content analysis showed virtually no differences between coating variants (Table 3).

**Table 3.**
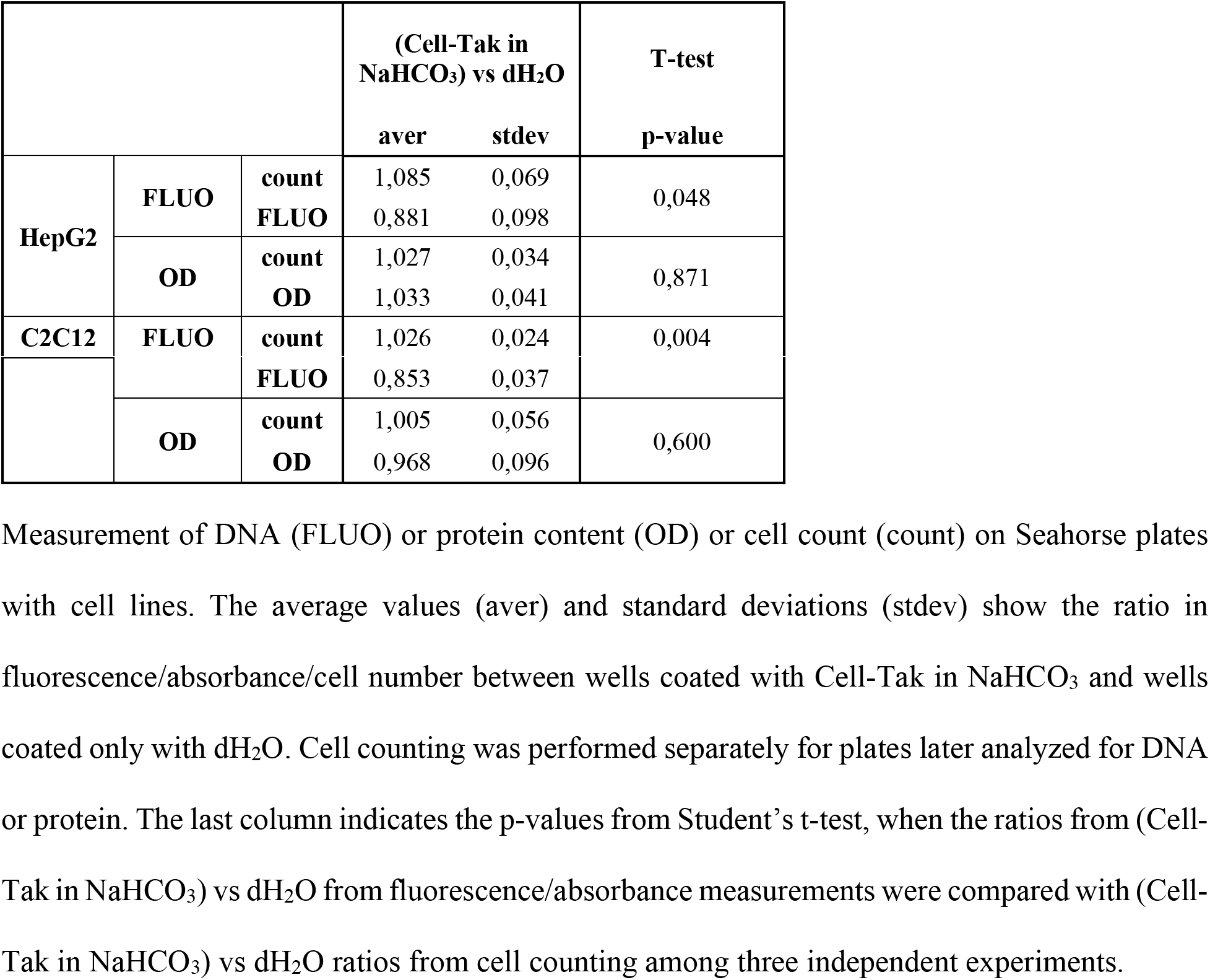
Results of DNA/protein/cell count deviations on Seahorse plates.

### Coating effects on metabolic flux normalization

In order to investigate the effect of coating in the intended context of metabolic flux measurements we used the standard mitochondrial stress test methodology consisting of measurements of basal cellular respiration, followed by measurements after the sequential additions of the ATP synthase inhibitor oligomycine, the uncoupler FCCP and a combination of mitochondrial complex I and III inhibitors rotenone and antimycine A. Prior to the experiments the appropriate concentration of FCCP to be used was established by titration for both cell lines. Only two types of coating were compared in these experiments: no coating (dH_2_O) or the commonly used Cell-Tak/NaHCO_3_.

We compared three normalization strategies (cell number, DNA content or protein content). When the results of the Seahorse measurement were normalized to cell number there were virtually no differences between plates coated with dH_2_O and Cell-Tak/NaHCO_3_. On the contrary, the normalization to DNA content showed a visible discrepancy between the coatings. When we normalized the OCR data to protein content very small differences were detected between coatings in both cell lines, somewhat more in C2C12 cells (Fig 1).

**Fig 1.**
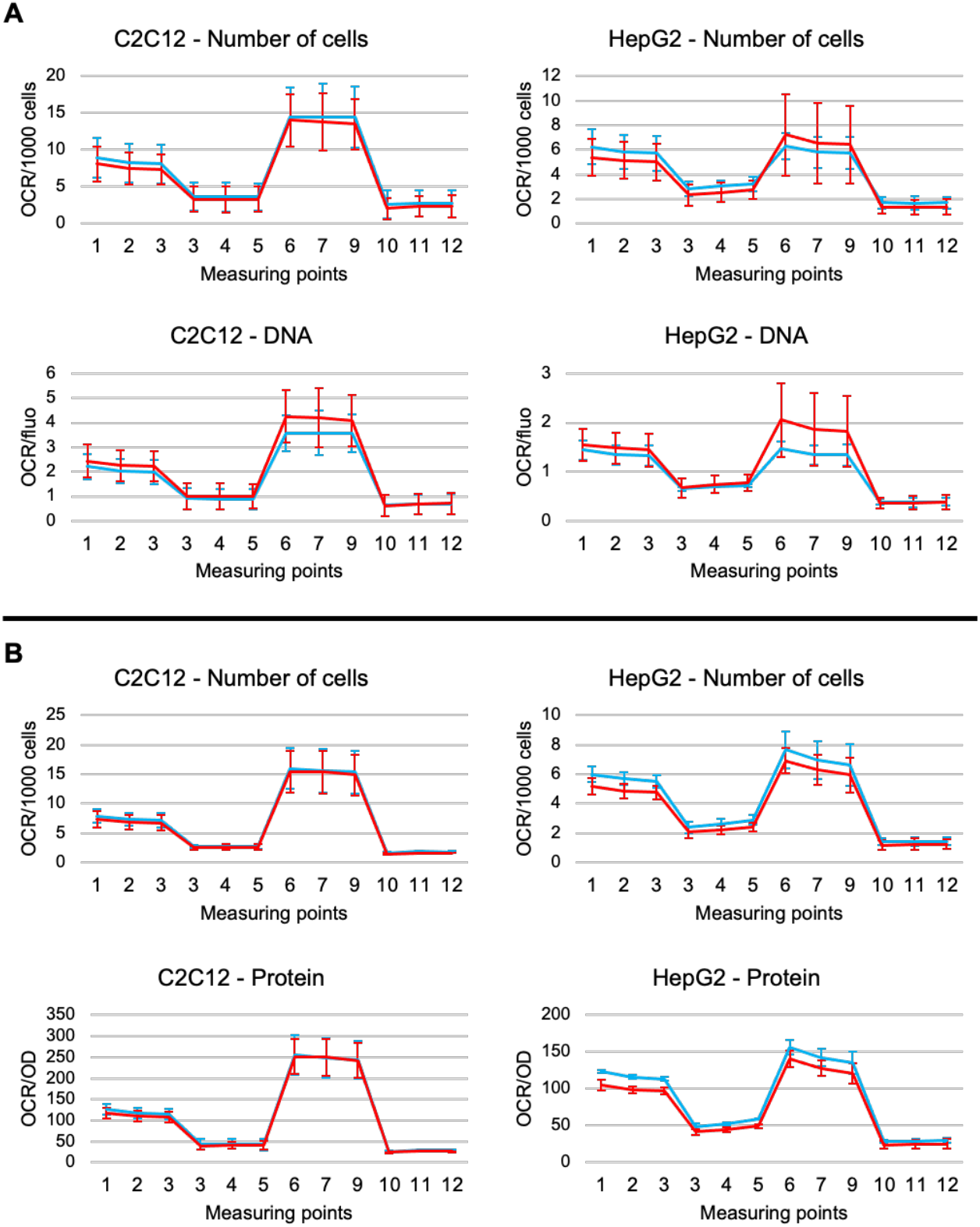
Normalized oxygen consumption rates in C2C12 and HepG2 cell lines differ due to coatings. **A** Data normalized to cell count and DNA content and **B** data normalized to cell count and protein content. Blue curves show uncoated wells (dH_2_O), red curves show wells coated with Cell-Tak/NaHCO_3_. Data shown as averages from three independent experiments +/- SEM. Asterisks indicate significant differences based on Student’s t-test (significance set at p<0.05).

### Coating effects on cell shape

During manual cell counting we noted that cells appear in two distinct cell shapes. A portion of cells retains a round form with defined and visible edges and the rest are considerably flatter without defined edges or with protrusions, presumable better attached to the plastic (Fig 2).

**Fig 2.**
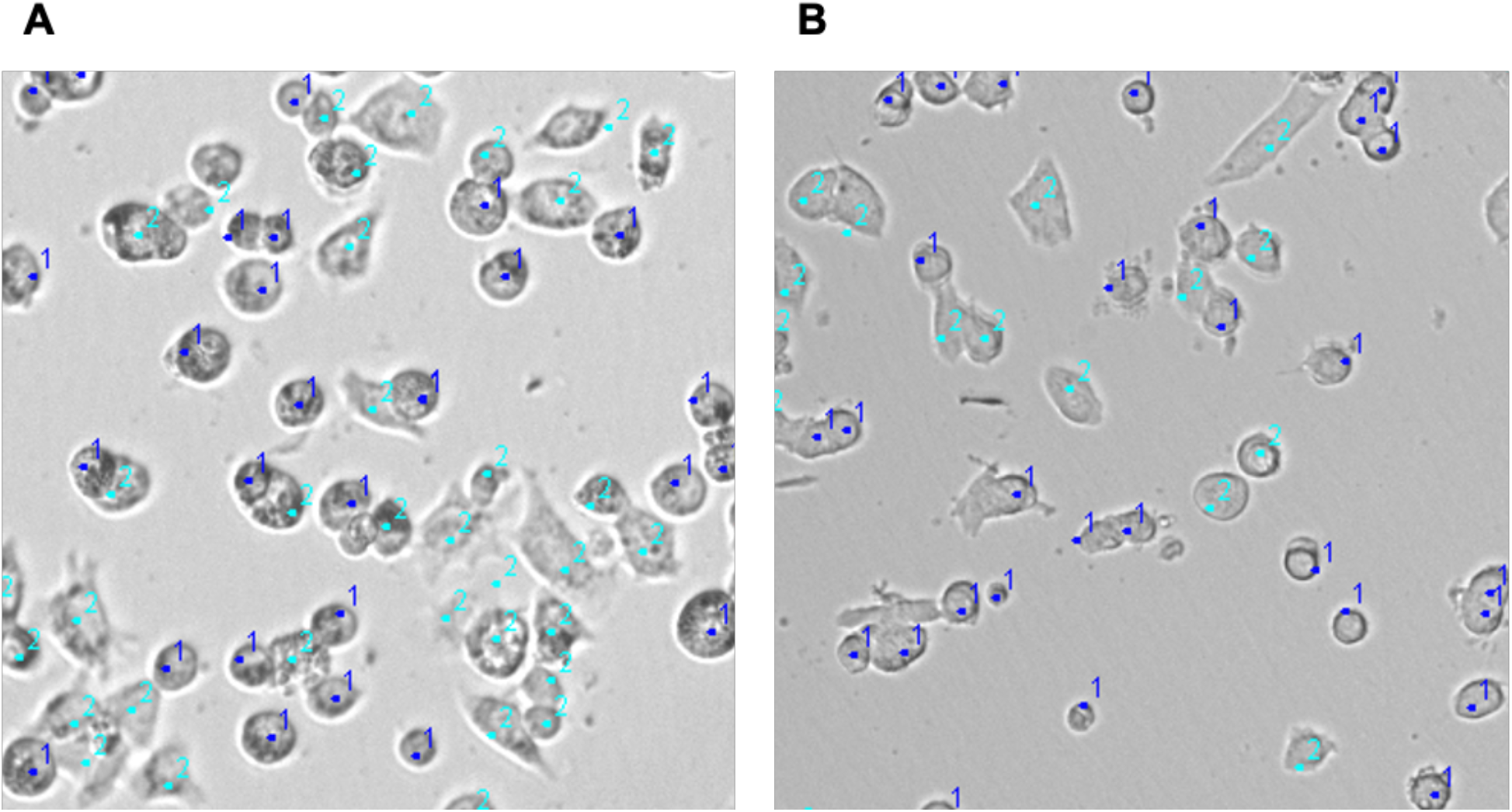
Cell shape variants identified during manual cell counting. In both cell lines (A: HepG2 and B: C2C12), two cell shape variants were detected in the Seahorse plates: round (1, dark blue) and flat (2, light blue).

The proportions of these cell shapes differed in coated vs. uncoated wells. In uncoated cells (dH_2_O), 88% of HepG2 cells were found in the round form, in wells coated with Cell-Tak/NaHCO_3_ the majority of cells were flat (62%). With C2C12 cells this difference was much less pronounced (Fig 3).

**Fig 3.**
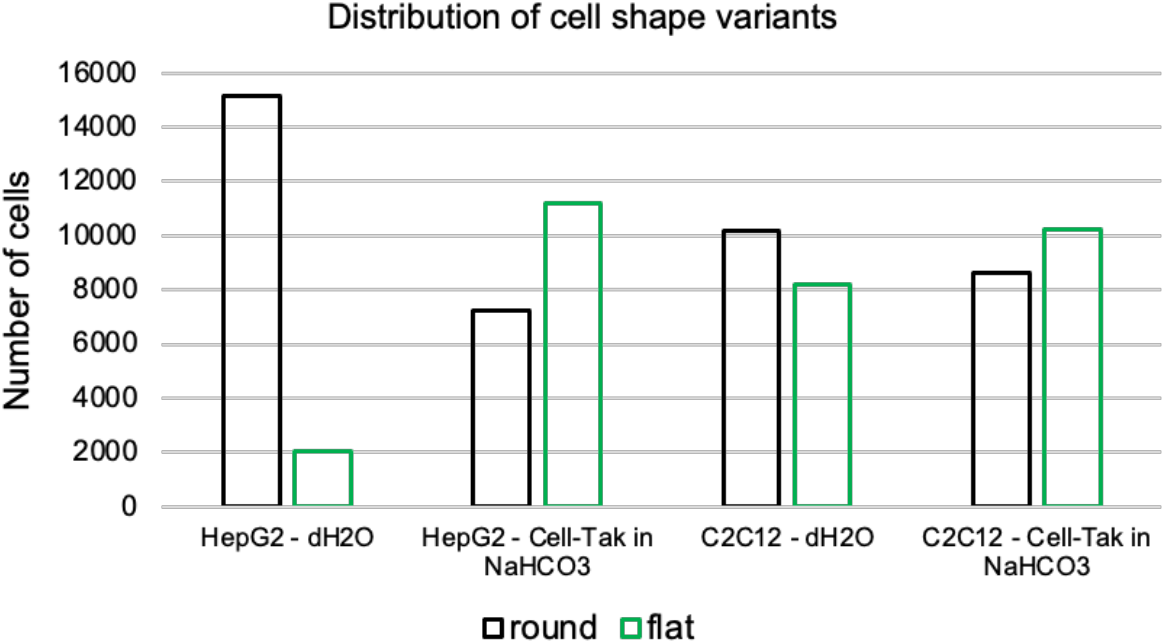
Differences in cell shapes influenced by plate coating type. The proportion of round/flat cells in uncoated wells (dH_2_O) and wells coated with Cell-Tak/NaHCO_3_. Data shown as averages from three independent experiments +/- SD.

## Discussion

In our previous experiments using the Seahorse extracellular flux analysis technology from Agilent we did not always get consistent data when we tried to normalize the metabolic measurements to DNA content using fluorescent dyes in wells coated with Cell-Tak. In this paper, we compared the effects of Cell-Tak coating on DNA and protein assays using standards as well as two cell lines, HepG2 and C2C12. Our results show that Cell-Tak coating affects fluorescent DNA assays, whether using pure DNA standards or in cultured cells. Wells coated with Cell-Tak/NaHCO_3_ had significantly decreased values of fluorescence compared to uncoated wells. A similar pattern was observed in cell lines cultured on coated vs. uncoated plastic but the effects appeared to be more cell line-specific and perhaps even specific to the type of plastic used as a different pattern was observed in normal tissue culture plates vs. Seahorse plates. In addition to differences in fluorescence values we also tended to observed much higher signal variability expressed in the standard deviation of measurements.

Total protein detection appears to be less affected by Cell-Tak coating. We observed virtually no difference when using a BSA standard, and an average 15% decrease in the measured protein content in both cell lines suggesting that things may be more complicated with total cellular protein content in a cell lysate. This difference in coating effects between DNA and protein assays could be due to the attachment of a portion of DNA molecules to the sticky coating, which may hinder the binding of the fluorescent reagent.

When we used all three normalization methods (protein, DNA, cell count) on real Seahorse extracellular flux data we saw a similar pattern as above but with some more cell line-specific observations. In the case of the murine myoblast cell line C2C12 normalizing to cell count or protein content produces virtually no difference between the OCR curves in uncoated vs. coated wells. Values normalized to DNA are, on the other hand, substantially higher in wells coated by Cell-Tak, which corresponds well with our observation of lower measured DNA content in coated wells.

In the case o HepG2 cells the picture is a little more complicated. While the discrepancy between uncoated and coated cells when normalized to DNA content is similar as with C2C12 cells, there also appears a slight increase in OCR in coated cells, which cannot be completely erased by the other normalization methods. This observation suggests a possible effect of Cell-Tak coating on the mitochondrial metabolism of these cells, which should be the topic of further investigations.

What may serve as a useful pointer for such further investigations is our observation that Cell-Tak coating affects cell shape. This effect on cell shape after attachment to the coated surface agrees with the previously published data about neuroblastoma cells [18] or hamster kidney cells and human histiocytic lymphoma cells [15]. Whether or not this change in cellular shape prevalence is connected to the observed OCR values remains to be investigated.

Our findings that the coating of cell culture or assay plastic with Cell-Tak may strongly influence DNA content measurements using PicoGreen fluorescence has important implications for the normalization of data from Seahorse extracellular flux analyses. One possible explanation for this is DNA molecules sticking to the coated surface. As this process is likely random, in addition to lower measured DNA fluorescence values we observed a very large variation in the data. Based on these results we suggest using cell count as the first-choice method and the total protein content as the second-choice technique for the normalization of Seahorse data whenever Cell-Tak coating is used. Researchers should work with caution when DNA fluorescence-based normalization strategies are utilized.

## Conflict of Interest

There is no conflict of interest.

## Funding

This study was supported by the Charles University institutional grant PROGRES Q36.

## Author contributions

JT, SM, and MS conceived the study, designed experiments, analyzed data and wrote manuscript. SM and MS performed the DNA and protein detection experiments and manual cell counting. AK, SM performed metabolic assay and JP the image analysis.

### Email addresses

MS

SM

AK

JP

JT

## Notes

### Competing Interest Statement

The authors have declared no competing interest.

